# Cell-intrinsic effects of TorsinA(ΔE) disrupt dopamine release in a mouse model of *DYT1-TOR1A* dystonia

**DOI:** 10.1101/2020.12.22.424006

**Authors:** Anthony M. Downs, Xueliang Fan, Radhika Kadakia, Yuping Donsante, H.A. Jinnah, Ellen J. Hess

## Abstract

*DYT1-TOR1A* dystonia is an inherited dystonia caused by a three base-pair deletion in the *TOR1A* gene (*TOR1A*ΔE). Although the mechanisms underlying the dystonic movements are largely unknown, abnormalities in striatal dopamine and acetylcholine neurotransmission are consistently implicated whereby dopamine release is reduced while cholinergic tone is increased. Because striatal cholinergic neurotransmission mediates dopamine release, it is not known if the dopamine release deficit is mediated indirectly by abnormal acetylcholine neurotransmission or if *Tor1a(ΔE)* acts directly within dopaminergic neurons to attenuate release. To dissect the microcircuit that governs the deficit in dopamine release, we conditionally expressed *Tor1a(ΔE)* in either dopamine neurons or cholinergic interneurons in mice and assessed striatal dopamine release using *ex vivo* fast scan cyclic voltammetry or dopamine efflux using *in vivo* microdialysis. Conditional expression of *Tor1a(ΔE)* in cholinergic neurons did not affect striatal dopamine release. In contrast, conditional expression of *Tor1a(ΔE)* in dopamine neurons reduced dopamine release to 50% of normal, which is comparable to the deficit in *Tor1a^+/ΔE^* knockin mice that express the mutation ubiquitously. Despite the deficit in dopamine release, we found that the *Tor1a(ΔE)* mutation does not cause obvious nerve terminal dysfunction as other presynaptic mechanisms, including electrical excitability, vesicle recycling/refilling, Ca^2+^ signaling, D2 dopamine autoreceptor function and GABA_B_ receptor function, are intact. Although the mechanistic link between *Tor1a(ΔE)* and dopamine release is unclear, these results clearly demonstrate that the defect in dopamine release is caused by the action of the *Tor1a(ΔE)* mutation within dopamine neurons.

## INTRODUCTION

Dystonia is a movement disorder characterized by abnormal muscle contractions that cause debilitating abnormal postures and/or twisting movements.^1^ *DYT1-TOR1A* dystonia, one of the most common inherited dystonias, is caused by a 3 base-pair in-frame deletion (Δgag) in the *TOR1A* gene that causes a single glutamic acid deletion in the torsinA protein, torsinA(ΔE).^2, 3^ While the causal mutation for *DYT1-TOR1A* dystonia has been known for some time, the functional link between the *TOR1A(ΔE)* mutation and dystonia remains unclear.

It is challenging to identify the neuroanatomical or cellular dysfunction that underlies dystonia because dystonia is not generally associated with a consistent neuropathology or neurodegeneration of specific nuclei. However, abnormalities in striatal dopamine (DA) and acetylcholine (ACh) neurotransmission are consistently implicated in multiple forms of dystonia.^4, 5^ Increased striatal cholinergic tone was initially suspected as an underlying pathophysiology in dystonia, because anticholinergics are often effective treatments.^6^ Indeed, abnormalities in striatal cholinergic neurotransmission are consistently observed in mouse models of dystonia,^5, 7–10^ including *DYT1-TOR1A* which exhibit an increase in striatal cholinergic tone.^11^ Reduced striatal DA neurotransmission is implicated in a diverse range of dystonias, including: idiopathic focal dystonias, secondary dystonia, and inherited dystonias, including *DYT1-TOR1A*.^12–22^ The *TOR1A* gene is abundantly expressed in DA neurons of the substantia nigra pars compacta (SNc) and the concentrations of DA and DA metabolites are abnormal in patients carrying the *TOR1A(ΔE)* mutation.^23–27^ Consistent with the abnormalities in DA handling observed in patients, striatal dopamine release is significantly reduced in mouse strains carrying the *Tor1a*(ΔE) mutation.^28–30^

Normal striatal function requires reciprocal interaction between DA and ACh neurotransmission.^31–33^ Under normal conditions, ACh released by striatal cholinergic interneurons (ChIs) activates nicotinic acetylcholine receptors located on DA terminals, which adjust the probability of DA release.^34–36^ DA released by nigrostriatal DA neurons activates D2 dopamine receptors expressed on ChIs to inhibit the firing rate of ChIs and reduce ACh release.^37^ In contrast, in mouse models of *DYT1-TOR1A*, D2 DA receptor activation enhances ChI firing rates. ^10, 11, 38, 39^ Further, an increase in extracellular striatal ACh has been observed in mouse models of *DYT1-TOR1A*, which has been postulated to cause the desensitization of nAChRs on DA terminals, resulting in the reduction in DA release observed in these mice.^11^ Because *Tor1a* is expressed in all cell types, ^25, 26, 40^ it is not known if the DA release deficit is caused indirectly by abnormal striatal ACh neurotransmission, or if it is the result of intrinsic effects of the *Tor1a(ΔE)* mutation in DA neurons. In this study, we used a genetic approach to conditionally express the *Tor1a(ΔE)* mutation in striatal ChIs or DA neurons to dissect the microcircuit that governs the abnormal DA release and interrogated potential mechanisms underlying the *Tor1a(ΔE)*-induced reduction in DA release. We found that expression of the *Tor1a(ΔE)* mutation in DA neurons alone was sufficient to recapitulate abnormalities in DA release while expression of the *Tor1a(ΔE)* mutation in ChIs alone did not affect DA release.

## MATERIALS AND METHODS

### Animals

For the conditional *Tor1a(ΔE)* expression experiments, *Tor1a^+/swap^* mice (*Tor1a^tm4.1Wtd^*/J; JAX #0208099) were bred with either heterozygous *DAT-cre* (JAX #006660) or *ChAT-cre* (JAX #031661) mice. The *Tor1a^swap^* allele carries a normal exon 5, which is flanked by loxP sites. Downstream of the normal exon 5 is another copy of exon 5, which carries the Δgag mutation.^41^ Without Cre recombinase expression, only the normal *Tor1a* gene is expressed. However, in the presence of Cre, the normal exon 5 is excised, which allows conditional expression of *Tor1a^ΔE^* in the endogenous *Tor1a* gene in a cell-type specific manner (**Fig 1**). For all other experiments, knockin mice heterozygous for the *Tor1a(ΔE)* mutation (*Tor1a^+/ΔE^*) and control littermates (*Tor1a^+/+^*) were used. All mice were inbred on C57BL/6J and were bred at Emory University. Animals were maintained on a 12 h light/dark cycle with *ad libitum* access to food and water. Mice were group housed with nestlets for environmental enrichment. Mice were genotyped using PCR with the following primers: *Tor1a^ΔE^* (forward primer, GCTATGGAAGCTCTAGTTGG; reverse primer CAGCCAGGGCTAAACAGAG); *Tor1a^swap^* (forward primer, TCCTCCCCCAAGTACATCAG; reverse primer CATAGCTCAGCCGTCCAGTC); *DAT-cre* (common primer, TGGCTGTTGGTGTAAAGTGG; WT reverse primer GGACAGGGACATGGTTGACT; cre reverse primer, CCAAAAGACGGCAATATGGT); *ChAT-cre* (WT forward primer, GCAAAGAGACCTCATCTGTGGA; cre forward primer, TTCACTGCATTCTAGTTGTGGT; common reverse primer, GATAGGGGAGCAGCACACAG). All studies were approved by the Institutional Animal Care and Use Committee at Emory University.

**Fig 1.**
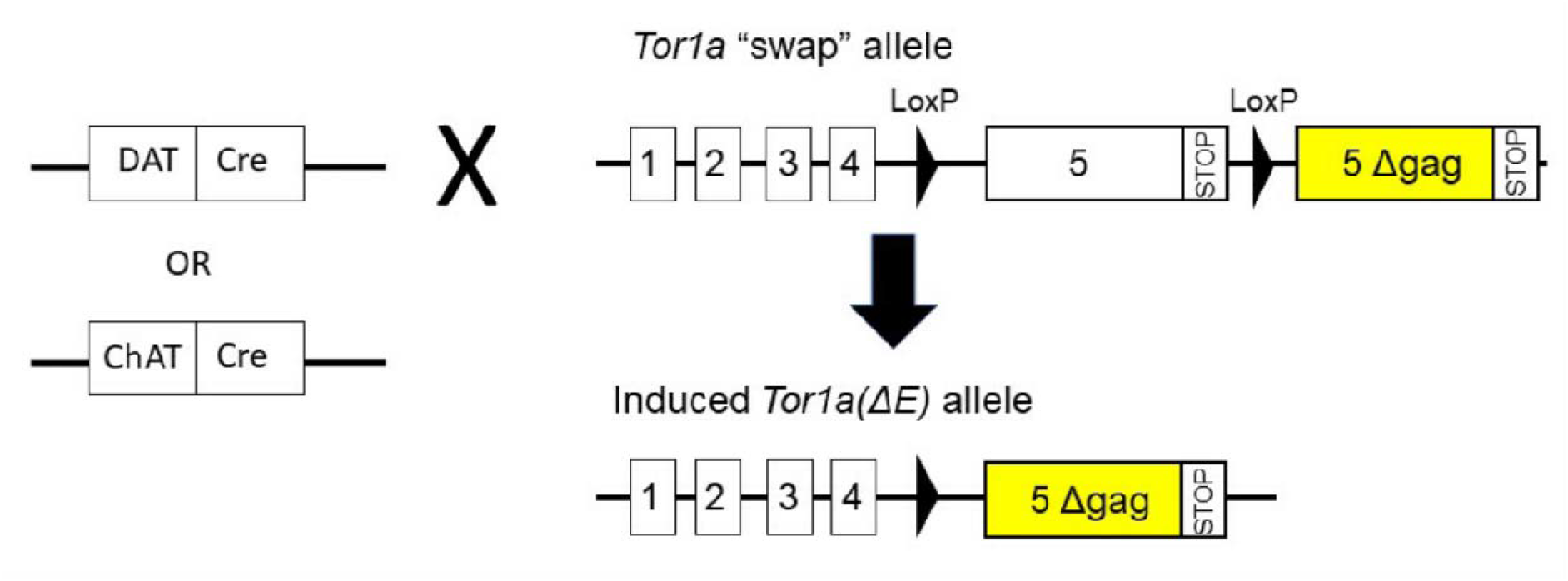
Schematic of cell-type specific *Tor1a(ΔE)* knockin approach. Mice carrying the “swa**p”** allele were crossed with either ChAT-cre or DAT-cre expressing mice to selective expre**ss** *Tor1a(ΔE)* in ChAT+ or DAT+ neurons respectively.

### Fast scan cyclic voltammetry

Fast scan cyclic voltammetry (FSCV) was performed as previously described.^29^ Mice were euthanized by cervical dislocation, and the brain was sectioned at 300μm using a vibratome in ice-cold, oxygenated sucrose artificial cerebral spinal fluid (aCSF) containing [in mM]: sucrose [194], NaCl [20], KCl [4.4], CaCl_2_ [1.2], MgCl_2_ [1.2], NaH_2_PO_4_ [1.2], NaHCO_3_ [25], D-glucose [11] at pH 7.4. Brain slices were collected in a holding chamber containing oxygenated, bicarbonate-buffered aCSF containing [in mM]: NaCl [126], KCl [2.45], CaCl_2_ [2.4], MgCl_2_ [1.2], NaH_2_PO_4_ [1.2], NaHCO_3_ [25], D-glucose [11] and maintained at room temperature for 45-60 min before experiments began.

All FSCV recordings were conducted in the dorsolateral striatum. This region was selected because it receives innervation from the motor cortex.^42^ A striatal slice at ~ Bregma +0.26 mm, where the anterior commissure is fused, was transferred to the recording chamber and perfused with oxygenated aCSF at 32°C. After a 30 min equilibration, a carbon fiber electrode was inserted approximately 50 μm into the surface of the slice and a bipolar tungsten stimulating electrode was placed approximately 200 μm away. DA release was evoked by either 1-pulse (600 μA, 4 ms pulse width) or 5-pulse 100 Hz electrical stimulation at 5 min inter-stimulus intervals to prevent rundown. The scan rate for voltammetry was 400 V/s from −0.4 V to 1.3 V to −0.4 V verses Ag/AgCl with a sampling rate of 10 Hz using a Chem-Clamp voltammeter-amperometer (Dagan Corporation, Minneapolis, MN, USA). FSCV experiments were conducted and analyzed using Demon voltammetry software (Wake Forest University).^43^ All drugs were diluted in aCSF and bath applied for 10-20 mins before recordings commenced to allow for equilibration. For the Ca^2+^ dose-response experiment, aCSF was prepared without calcium. Ca^2+^ was then added from a 1M CaCl_2_ stock solution to prepare the appropriate Ca^2+^ concentration. All electrodes were calibrated to known DA standards in aCSF using a custom-made flow cell.

### Tissue monoamines

Mice were euthanized by cervical dislocation and striata were rapidly dissected, frozen on dry ice, and stored at −80 °C. For high performance liquid chromatography (HPLC) analysis of monoamines, tissue was homogenized in 100mM perchloric acid by probe sonication at 4°C and centrifuged at 10,000 g for 10 min. The supernatant was collected and filtered through 0.45 μm PVDF membranes prior to analysis. The cell pellet was solubilized in 2% SDS and protein concentrations were determined using a BCA assay (ThermoFisher, Waltham, MA, USA). Monoamines were measured using HPLC with electrochemical detection as previously described.^44^ The HPLC system consisted of an ESAS MD-150 x 3.2 mm column, an ESA 5020 guard cell, and an ESA 5600A Coularray detector with an ESA 6210 detector cell (ESA, Bedford MA). The guard cell potential was 475 mV; and the analytical cell potentials were 175, 100,350, and 450 mV. Samples were eluted at a flow rate of 0.4 mL/min with a mobile phase composed of [in mM], [1.7] 1-octanesulfonic acid sodium, [75] NaH_2_PO_4_, 0.25% triethylamine, and 8% acetonitrile at pH 2.9. Monoamines were identified by both retention time and electrochemical profile and compared with known standards.

### No-net flux microdialysis

Microdialysis was performed as previously described.^45^ Briefly, concentric microdialysis probes were manufactured in-house and calibrated with 100 ng/mL DA in aCSF containing [in mM]: NaCl [147], KCl [3.5], CaCl_2_ [1.2], MgCl_2_ [1.2], NaH_2_PO_4_ [1] at pH 7.0-7.4. Mice were anesthetized with isoflurane and the probe was implanted in the dorsal striatum (from bregma: anterior 0.6 mm, lateral 1.7 mm, and ventral 4.5 mm). The probe was perfused with aCSF at a flow rate of 0.6 μL/min while the mice habituated to the experimental chamber overnight. The following morning, the probe was perfused with aCSF and 0, 2, 10 or 20 nM DA (DA_in_) presented in pseudo-random order. After a 30 min equilibration period for each concentration of DA, 3 samples (40 min each) were collected (DA_out_). Samples were stored at −80 °C until HPLC analysis as described above. Extracellular DA levels were determined using simple linear regression analysis of the gain and loss of DA in the dialysate (DA_in_-DA_out_) versus DA_in_. After sample collection, the probe location was verified by reverse dialysis of 3% bromophenol blue, and only mice with a probe correctly located in the dorsal striatum were included in the analysis.

### Immunohistochemistry

Mice were euthanized with isoflurane, transcardially perfused with ice-cold normal saline followed by perfusion with ice-cold 4% paraformaldehyde in phosphate-buffered saline (PBS). Brains were removed and placed in 4% paraformaldehyde in PBS overnight at 4°C. Brains were then serially incubated in 20% and 30% sucrose solutions (w/v) for 24 hours each. Sections were cut at 30 μm using a freezing microtome and stored in cryoprotectant at −20 °C.

Immunostaining for choline acetyltransferase (ChAT) or tyrosine hydroxylase (TH) was performed in free-floating sections were processed using the avidin-biotin complex method (ABC) (Vector Laboratories, Burlingame, CA, USA). Sections were washed in tris-buffered saline (TBS) and permeabilized with 0.4% Triton-X100 in TBS with 0.5% bovine serum albumin (BSA) for 10 mins. Sections were then incubated with 1% H_2_O_2_ for 30 mins to eliminate endogenous peroxidase activity. Sections were incubated in primary antibody using goat anti-ChAT 1:500 (Millipore cat # AB144P) or rabbit anti-TH 1:1000 (Pel-Freez cat # P4010-0) diluted in TBS containing 0.4% Triton-X100, 1% BSA and 5% normal donkey serum for 16 hours at 4°C with gentle agitation. Sections were then washed and incubated with biotinylated secondary antibody for 2 h and then incubated with the avidin-biotin complex reagent solution for 1 h. Chromogen was developed using the ImmPACT VIP peroxidase substrate kit (Vector). Sections were then mounted on charged slides and coverslipped.

For TH and TorsinA double-labeling, free-floating sections were washed in TBS and permeabilized with 0.4% Triton-X100 in TBS with 0.5% BSA for 10 mins. Sections were blocked in TBS containing 0.4% Triton-X100, 1% BSA and 5% normal donkey serum for 2 hours at RT with gentle agitation. Rabbit anti-TorsinA 1:250 (Abcam cat # ab34540) and mouse anti-TH 1:1000 (Immunostar cat # 22941) were diluted in the same blocking solution and incubated for 16 hr at 4 °C with gentle agitation. Sections were then washed in TBS, permeabilized, blocked in 5% normal donkey serum, and incubated with Alexa Fluor secondary antibodies (Invitrogen, Carlsbad, CA, USA) at 1:200 dilution for 2 hrs. Sections were then washed again in TBS, mounted on charged slides, and coversliped.

### Laser capture microdissection

Mice were euthanized by cervical dislocation, brains were rapidly removed, embedded in optimal cutting temperature solution (Sakura Fientek, Torrance, CA, USA) and frozen in a dry ice/isopentane slurry. Sections were cut at 10 μm using a cryostat and sections containing the striatum and SNc/ventral tegmental area (VTA) were collected on Leica PET FrameSlides (Leica Microsystems, Buffalo Grove, IL, USA). Slides were stored at −80 °C until staining.

Immunostaining for ChAT or TH was performed using the following antibodies: rabbit anti-TH 1:200 (Pel-Freez cat # P4010-0) or goat anti-ChAT 1:25 (Millipore cat # AB144P). Sections were fixed in an acetone/methanol mixture (1:1) at −20 °C for 15 mins. Slides were washed in diethyl pyrocarbonate (DEPC)-treated water before incubation with primary antibody for 20 min at RT. Slides were then washed with DEPC-treated PBS before incubation with Alexa 488-conjugated secondary antibody for 5 min at RT. Slides where then washed in DEPC-treated PBS, treated with DAPI for 2 mins, and then washed with DEPC-treated PBS again. Slides were then dehydrated in graded ethanols and xylenes and then air dried. Slides proceeded immediately to laser capture microdissection (LCM). LCM was performed using an Arcturus XT Laser Capture Microdissection system (Thermo Fisher) with a Nikon Eclipse Ti-E microscope base (Nikon Instruments, Melville, NY, USA). Microdissection was performed using both an infrared laser and ultraviolet light. Samples were collected on CapSure HS LCM Caps (Thermo Fisher).

### RNA isolation and sequencing

RNA isolation and purification were performed using the PicoPure RNA Isolation kit (Thermo Fisher). Briefly, a CapSure HS LCM cap containing microdissected cells was applied to a 0.5 mL microcentrifuge tube containing 10 μL extraction buffer and incubated for 30 min at 42 °C. 10 μL of 70% ethanol was then added and mixed thoroughly. RNA was then purified using a nucleic acid binding column with on-column DNase treatment (RNase-free DNase set, QIAGEN, Germantown, MD, USA). RNA was then eluted from the column in elution buffer.

RNA was reverse transcribed using LunaScript RT SuperMix Kit (New England Biolabs, Ipswich, MA, USA). To confirm that striatal ChIs or midbrain dopaminergic neurons were collected via LCM, RT-qPCR was performed using Luna Universal qPCR master mix (New England Biolabs) with primers specific to 18s rRNA (forward primer 5’-GGACCAGAGCGAAAGCATTTGCC-3’ and reverse primer 5’-

TCAATCTCGGGTGGCTGAACGC-3’), ChAT (forward primer 5’-GTGAGACCCTGCAGGAAAAG-3’ and reverse primer 5’-GCCAGGCGGTTGTTTAGATA-3’), or TH (forward primer 5’-GGTATACGCCACGCTGAAGG-3’ and reverse primer 5’-TAGCCACAGTACCGTTCCAGA-3’). To verify cell-type specific recombination of the *Tor1a^+/swap^* allele, cDNA encompassing the *Tor1a(Δgag)* mutation was amplified using forward primer 5’-GCCGTGTCGGTCTTCAATAA-3’ and reverse primer 5’-ACAGTCTTGCAGCCCTTGTC-3’. The PCR product was purified using QIAquick PCR purification kit (QIAGEN) and sequenced using the primer 5’-GCCGTGTCGGTCTTCAATAA-3’. This primer is located within exon 4 of the *Tor1a* gene to ensure that cDNA derived from mRNA, but not genomic DNA was sequenced. Sequencing chromatographs were analyzed using 4Peaks software (Mekentosj, Amsterdam, NL).

### Image collection

Slides were imaged under bright field illumination using an Olympus BX51 camera equipped with an Olympus DP72 camera and Olympus CellSens software. Immunofluorescence was imaged using a Leica SP8 confocal microscope using Leica Acquisition Suite (LAX S) (Leica Microsystems). Images were taken from fields including the SNc or medial forebrain bundle using a 40X objective and an optical thickness of ~1 μm. The same optical settings were used for image acquisition in each region from each mouse.

### Statistical analysis

Data are presented as means with standard error. Dose responses were analyzed with non-linear regression to determine EC_50_ or IC_50_. IC_50_ and EC_50_ were analyzed using two-tailed Student’s *t*-test. All other data were analyzed using one-way ANOVA with *post hoc* Dunnett’s multiple comparison test or two-way ANOVA with Sidak’s multiple comparisons test. All analyses were performed using Graphpad Prism 8 (https://www.graphpad.com/). Statistical significance was defined as \**p* < 0.05, \*\**p* < 0.01, \*\*\**p* < 0.001.

### Compounds

Quinpirole and CGP 88584 were purchased from Tocris (Minneapolis, MN, USA).

### Data availability

The data that support the findings of this study are available from the corresponding author, upon reasonable request.

## RESULTS

### Conditional expression of *Torla(ΔE)* in striatal ChIs does not affect striatal DA release

Because ChI dysfunction is implicated in *DYT1-TOR1A* dystonia and cholinergic neurotransmission plays a critical role in regulating DA release,^5, 7, 8, 34, 46^ we hypothesized that expression of *Tor1a(ΔE)* in striatal ChIs mediates the deficit in striatal DA release. To test this hypothesis, we generated mice that were heterozygous knockin for the *Tor1a^+/ΔE^* mutation only in cholinergic neurons while other cells harbored the normal (*Tor1a^+/+^*) alleles. To produce this cell-type specific genocopy of the human disease-causing genotype (*TOR1A^+/^*^ΔE^), we bred ChAT-cre mice, which drives Cre recombinase only in neurons that also express the choline acetyltransferase gene, onto the *Tor1a^swap^* mouse strain (*ChAT-cre; Tor1a^+/swap^* mice). To validate conditional expression of *Tor1a(ΔE)* in only ChIs, we used LCM to dissect ChAT-positive ChIs and sequenced the *Tor1a* gene from cDNA derived from these cells. The sequencing demonstrated heterozygous expression of *Tor1a(ΔE)* in ChIs but not ChAT-negative cells from the striatum in *ChAT-cre; Tor1a^+/swap^* mice nor in ChIs collected from *ChAT-cre; Tor1a^+/+^* mice (**Fig 2A**). As additional confirmation that ChIs were collected by LCM, we performed RT-qPCR for ChAT and found significant enrichment of ChAT transcripts in putative ChIs compared to other LCM-captured striatal cells where no ChAT immunostaining was detected (data not shown). Further, ChAT immunostaining in the dorsolateral CPu of *ChAT-cre; Tor1a^+/swap^* mice was indistinguishable from *ChAT-cre; Tor1a^+/+^* mice (**Fig 2B**).

**Fig 2.**
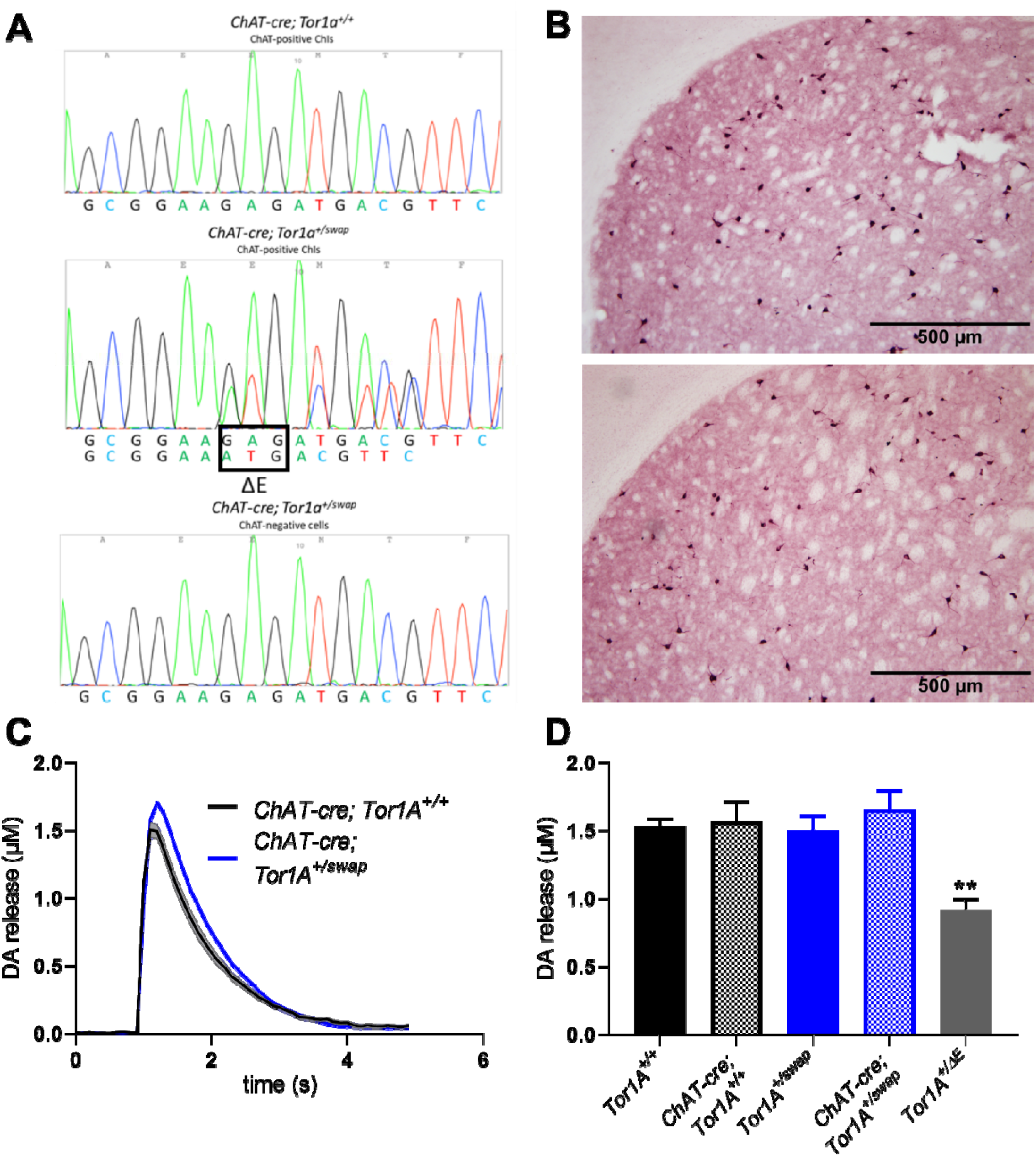
Expression of *Tor1a(ΔE)* in ChAT+ neurons does not alter DA release. (**A**) Validation of cell-type specific expression of *Tor1a(ΔE)* in striatal cholinergic cells. Sequencing traces demonstrate that *ChAT-cre; Tor1a^+/swap^* mice express one copy of the *Tor1a(ΔE)* mutation in cholinergic cells but not ChAT-negative cells in the striatum. Cholinergic cells from *ChAT-cre*; *Tor1a^+/+^* do not express *Tor1a(ΔE)*. (**B**) Immunostaining for ChAT in the striatum. There is no obvious change in the number or distribution of striatal ChIs in either *ChAT-cre; Tor1a^+/+^* (upper figure) or *ChAT-cre; Tor1a^+/swap^* mice (lower figure). (**C**) Striatal DA release is unchanged in *ChAT-cre; Tor1a^+/swap^*; versus *ChAT-cre; Tor1a^+/+^* (*n*= 5). (**D**) Striatal DA release is significantly reduced in *Tor1a^+/ΔE^* versus all other genotypes (*n* = 5 for all except *Tor1a^+/ΔE^, n* = 17) (Values represent mean ± SEM).

We assessed total tissue striatal monoamine concentrations in *ChAT-cre; Tor1a^+/swap^* mice and control genotypes including *Tor1a^+/swap^* and *ChAT-cre; Tor1a^+/+^* to determine if expression of *Tor1a(ΔE)* in cholinergic neurons alone affected dopamine neurochemistry.

Consistent with our previous work demonstrating that DA and DA metabolite concentrations are normal in *Tor1a^+/ΔE^* knockin mice,^45^ we found that there were no significant differences in the striatal concentrations of DA (one-way ANOVA, F_2,14_ = 0.5173, p = 0.61), DOPAC (F_2,14_ = 0.04246, 0.96), HVA (F_2,14_ = 0.4396, p = 0.65), or 3-MT (F_2,14_ = 0.6304, p = 0.63), among *Tor1a^+/swap^, ChAT-cre; Tor1a^+/+^*, or *ChAT-cre; Tor1a^+/swap^* mice (**Table 1**).

**Table 1:** Tissue monoamines in the striatum. Results are expressed as average ng/mg tissue. Abbreviations: DOPAC (3,4-dihydroxyphenylacetic acid), 3-MT (3-methoxytyramine), and HVA (homovanillic acid)

|  | *Tor1A*<sup>+/-swap</sup> | *DAT-cre*;<br>*Tor1A*<sup>+/+</sup> | *DAT-cre*;<br>*Tor1A*<sup>+/-swap</sup> | *ChAT-cre*;<br>*Tor1A*<sup>+/+</sup> | *ChAT-cre*;<br>*Tor1A*<sup>+/-swap</sup> |
| --- | --- | --- | --- | --- | --- |
| DA | 37.01±4.69 | 29.15±3.07 | 37.75±5.60 | 42.84±4.05 | 43.56±5.85 |
| DOPAC | 1.96±0.31 | 1.71±0.34 | 2.02±0.39 | 1.85±0.12 | 1.93±0.32 |
| HVA | 3.20±0.39 | 3.32±0.34 | 4.02±0.91 | 3.77±0.30 | 3.70±0.62 |
| 3-MT | 1.73±0.20 | 1.71±0.19 | 2.33±0.46 | 1.97±0.23 | 2.03±0.27 |

To determine if expression of the disease-causing genotype in cholinergic neurons was sufficient to induce a deficit in DA release, FSCV was used to measure evoked DA release in the dorsal striatum. DA was identified by its characteristic oxidation peak at +600mV and reduction peak at −200mV (data not shown). There was no significant difference between the kinetics of DA uptake *ChAT-cre; Tor1a^+/+^* and *ChAT-cre; Tor1a^+/swap^* based on tau (*ChAT-cre; Tor1a^+/+^* 0.73 ± 0.03 s; *ChAT-cre; Tor1a^+/swap^* 0.78 ± 0.07 s, Student’s *t* test, p = 0.57) (**Fig 2C**). Similarly, there was no significant difference in peak DA release between *ChAT-cre; Tor1a^+/swap^* and the control genotypes *Tor1a^+/+^*, *Tor1a^+/swap^* and *ChAT-cre; Tor1a^+/+^* (**Fig 2D**). As we have shown previously, *Tor1a^+/ΔE^* knockin mice, which we used as a positive control, exhibited a significant reduction in DA release compared to *Tor1a^+/+^* mice. DA release in *Tor1a^+/ΔE^* knockin mice was also significantly reduced compared to *ChAT-cre; Tor1a^+/swap^ (Tor1a^+/ΔE^*, 0.91 ± 0.08 μM DA; *ChAT-cre; Tor1a^+/swap^*, 1.66 ± 0.14 μM DA, F_4,32_ = 10.54, *p* < 0.0001; Dunnet’s multiple comparisons test, *p* = 0.0013) and all other genotypes. We also examined *in vivo* extracellular striatal DA after conditional expression of *Tor1a(ΔE)* in ChIs using no-net flux microdialysis. There was no significant difference in extracellular DA concentrations between *ChAT-cre; Tor1a^+/+^* and *ChAT-cre; Tor1a^+/swap^ (ChAT-cre; Tor1a^+/+^* 3.15 ± 0.64 nM DA; *ChAT-cre; Tor1a^+/swap^* 3.07 ± 0.51 nM DA, Student’s *t* test, p = 0.93).

### Conditional expression of *Tor1a(ΔE)* in DA neurons causes abnormal DA release

Because expression of the *Tor1a(ΔE)* mutation in cholinergic neurons had no effect on striatal DA release, we next hypothesized that the *Tor1a(ΔE)* mutation acts intrinsically within DA neurons to cause the deficit in DA release. To test this, DAT-cre, which drives Cre recombinase only in neurons that also express the DA transporter, was bred onto the *Tor1a^swap^* mouse strain (*DAT-cre; Tor1a^+/swap^* mice). To validate conditional expression of *Tor1a(ΔE)* in only midbrain DA neurons, we used LCM to dissect TH-positive DA neurons and sequenced the *Tor1a* gene from cDNA derived from these cells. The sequencing demonstrated heterozygous expression of *Tor1a(ΔE)* in TH-positive neurons from *DAT-cre; Tor1a^+/swap^* mice but TH-negative cells and TH-positive DA neurons collected from *DAT-cre; Tor1a^+/+^* mice expressed only the normal *Tor1a* transcript (**Fig 3A**). We further validated that DA neurons were collected by using RT-qPCR for TH and found significant enrichment of TH transcripts in putative DA neurons versus other cells (data not shown). Further, TH immunostaining in midbrain DA neurons in *DAT-cre; Tor1a^+/swap^* mice was indistinguishable from *DAT-cre; Tor1a^+/+^* mice (**Fig 3B**).

**Fig 3.**
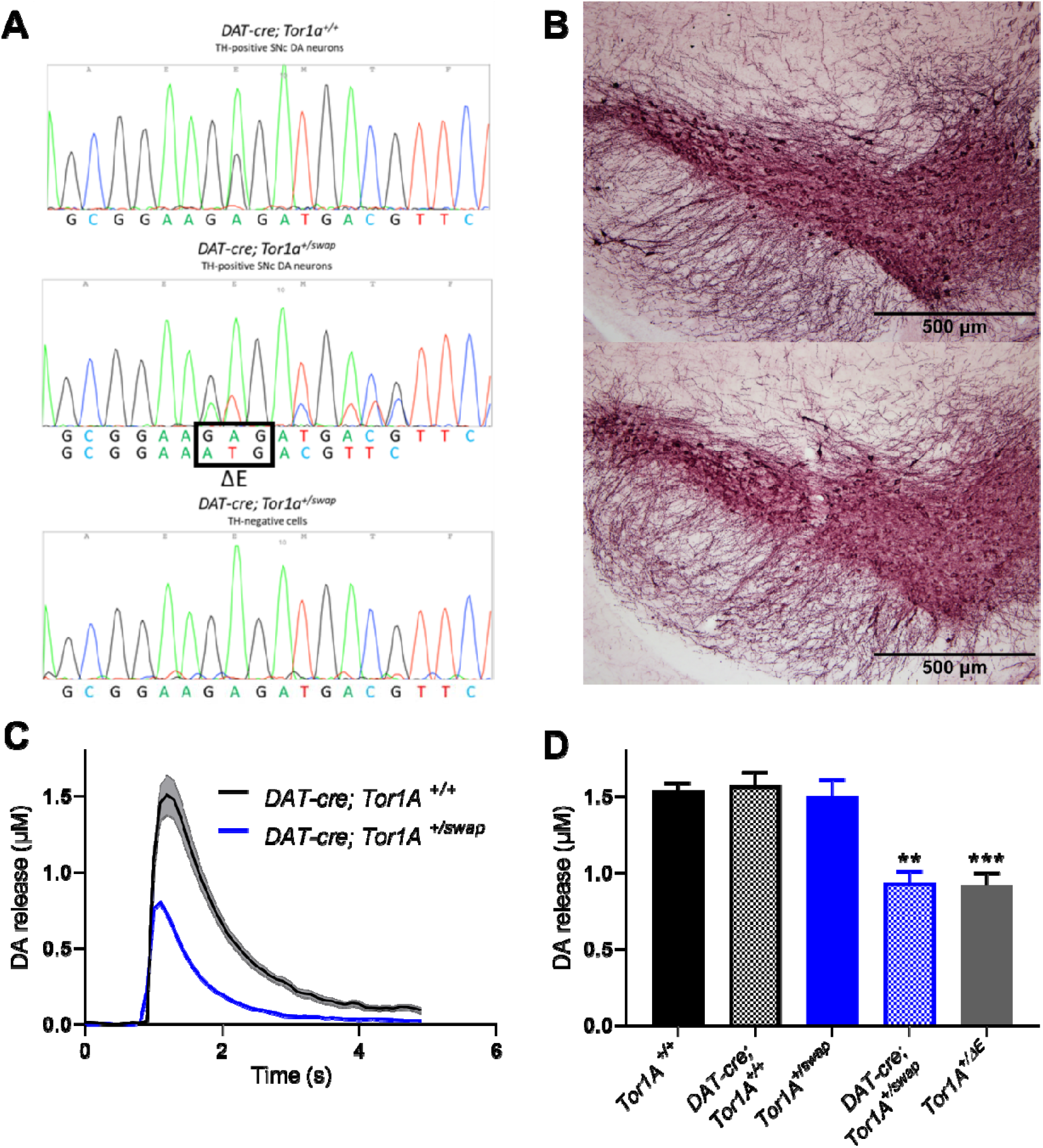
Expression of *Tor1a(ΔE)* in DA neurons reduces striatal DA release. (**A**) Validation of cell-type specific expression of *Tor1a(ΔE)* in SNc DA neurons. Sequencing traces demonstrate that *DAT-cre; Tor1a^+/swap^* mice express one copy of the *Tor1a(ΔE)* mutation in SNc DA neuro**ns** but not TH-negative cells in the midbrain. SNc DA neurons from *DAT-cre; Tor1a^+/+^* do not express *Tor1a(ΔE)*. (**B**) Immunostaining for TH in the SNc. There is no obvious change in the number or distribution of SNc DA neurons in either *DAT-cre; Tor1a^+/+^* (upper figure) or *DAT-cre; Tor1a^+/swap^* mice (lower figure). (**C**) Striatal DA release is significantly reduced in *DAT-cre; Tor1a^+/swap^* versus *DAT-cre; Tor1a^+/+^* (*n*= 5). (**D**) Striatal DA release is reduced in both *DAT-cre; Tor1a^+/swap^* and Tor1a^+/ΔE^ versus all control genotypes. DA release is not significantly different between *DAT-cre; Tor1a^+/swap^* and *Tor1a^+/ΔE^* (Values represent mean ± SEM).

To determine if conditional expression of the *Tor1a*(ΔE) mutation in DA neurons affected DA neurochemistry, we assessed total striatal monoamine concentrations. There were no significant differences in the striatal concentrations of DA (One-way ANOVA, F_2,15_ = 0.8622, p = 0.44), DOPAC (F_2,15_ = 0.2256, p = 0.80), HVA (F_2,15_ = 0.5351, p = 0.53), or 3-MT (F_2,15_ = 1.315, p = 0.30) among *Tor1a^+/swap^, DAT-cre; Tor1a^+/+^*, or *DAT-cre; Tor1a^+/swap^* mice (**Table 1**).

We then used FSCV to measure evoked DA release in the dorsolateral striatum after conditional expression of *Tor1a(ΔE)* in DA neurons. There was a significant reduction in peak DA release in *DAT-cre; Tor1a^+/swap^* relative to *DAT-cre; Tor1a^+/+^* and *Tor1a^+/swap^* (One-way ANOVA, F_4,32_ = 11.24, p<0.0001; Dunnett’s *post hoc* test, p = 0.008) (**Fig 3C**). There was no change in the kinetics of DA uptake between *DAT-cre; Tor1a^+/+^* and *DAT-cre; Tor1a^+/swap^ (DAT-cre; Tor1a^+/+^* 0.78 ± 0.06 s; *DAT-cre; Tor1a^+/swap^* 0.67 ± 0.08, Student’s *t* test, p = 0.33). The decrement in DA release in *DAT-cre; Tor1a^+/swap^* mice, which express the mutation in only DA neurons, was comparable to that of *Tor1a^+/ΔE^* knockin mice, which express the mutation in all cell types (*DAT-cre; Tor1a^+/swap^*, 0.94 ± 0.07 μM DA; *Tor1a^+/ΔE^*, 0.91 ± 0.08 μM DA, Dunnet’s multiple comparison’s test, p = 0.99) (**Fig 3D)** We also examined striatal extracellular DA concentrations *in vivo* in awake behaving *DAT-cre; Tor1a^+/swap^* and control mice using nonet-flux microdialysis. Extracellular DA was significantly reduced in *DAT-cre; Tor1a^+/swap^* versus *DAT-cre; Tor1a^+/+^ (DAT-cre; Tor1a^+/+^* 5.36 ± 0.03 nM DA; *DAT-cre; Tor1a^+/swap^* 2.90 ± 0.55 nM DA, Student’s *t* test, p = 0.048). Overall, these results demonstrate that expression of the *Tor1a(ΔE)* in DA neurons causes a reduction in DA release without affecting DA neurochemistry.

### Effects of the *Tor1a(ΔE)* mutation on the excitability of DA terminals

Our results suggest that the decrement in DA release is caused by intrinsic effects of the *Tor1a(ΔE)* on midbrain DA neurons, but the mechanisms underlying the DA release defect are unknown. To determine if abnormal excitability could account for the deficit in release, we conducted a current-response experiment using FSCV. Stimulation current was varied between 200 μA and 800 μA and peak DA release was measured relative to our standard paradigm of 600 μA stimulation current. The EC_50_ was comparable between control and *Tor1a^+/ΔE^* knockin mice (EC_50_: *Tor1a^+/+^* = 405.5 ± 8.73 μA; *Tor1a^+/ΔE^* = 405.3 ± 6.11 μA; Student’s *t*-test, p = 0.99, n=3; **Fig 4A**). Further, when the data was expressed as raw [DA], DA release in *Tor1a^+/ΔE^* was reduced at all stimulation currents tested, except for the lowest stimulation currents between 200-300 μA which produced no DA release, and even the highest stimulation current did not overcome the deficit in DA release observed in *Tor1a^+/ΔE^* knockin mice (**Fig 4B**).

**Fig 4:**
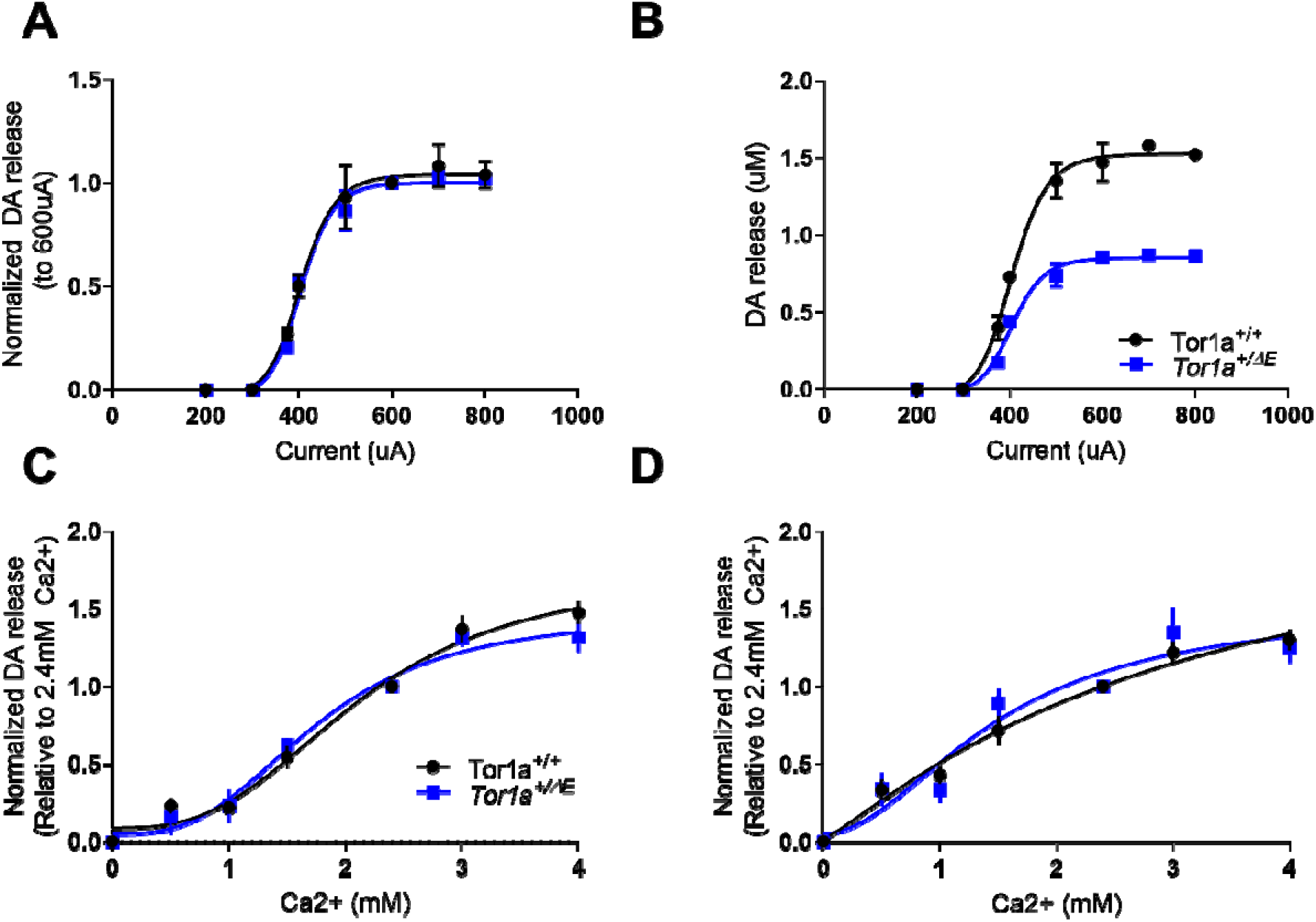
Sensitivity of striatal DA release to increasing stimulation current and extracellular Ca^+^. (**A**) There is no difference in the sensitivity of DA release to electrical stimulation of increasing current. There is no significant difference in EC_50_ between control and *Tor1a^+/ΔE^* mice (*n* = 3). (**B**) Increasing stimulation current does not normalize DA release in *Tor1a^+/ΔE^* mice to control levels (*n* = 3). (**C**) Control and *Tor1a^+/ΔE^* mice have similar sensitivity to varyi**ng** concentrations of Ca^2+^ after 1-pulse electrical stimulation. There was no significant difference in EC_50_ between control and *Tor1a^+/ΔE^* (*n* = 6). (**D**) Control and *Tor1a^+/ΔE^* mice have similar sensitivity to varying concentrations of Ca^+^ after 5-pulse 100 hz electrical stimulation. There was no significant difference in EC_50_ between control and *Tor1a^+/ΔE^* mice (*n* = 6). Values represent mean ± SEM.

### Sensitivity to extracellular Ca^2+^ concentration in *Tor1a^+/ΔE^* mice

Previous studies have shown that striatal DA release is highly sensitive to extracellular Ca^2+^ concentration and mediated by Ca_V_2.2 channels^47^. Further, abnormal Ca_V_2.2 Ca^2+^ channel activation has been observed in a transgenic mouse model carrying the *Tor1a(ΔE)* mutation suggesting that the regulation of release by Ca^2+^ is abnormal in *Tor1a^+/ΔE^* mice.^10^ To test this hypothesis, we assessed DA release in response to varying concentrations of Ca^2+^ (0-4mM Ca^2+^). There was no difference in the sensitivity to Ca^2+^ between genotypes with 1-pulse stimulation (**Fig 4C**) (EC_50_: *Tor1a^+/ +^* = 2.12 ± 0.21 mM; *Tor1a^+/ΔE^* = 1.75 ± 0.36 mM; Student’s *t*-test, p = 0.056, n=6). There was also no significant difference in the Hill slope between the genotypes indicating there is no alteration in cooperativity for Ca^2+^ (Hill coefficient: *Tor1a^+/ +^* = 2.92 ± 0.60; *Tor1a^+/ΔE^* = 2.95 ± 0.76; Student’s *t*-test, p = 0.98 n=6). We also tested the effect of altered Ca^2+^ concentration on 5-pulse 100 hz stimulations, because experimental manipulations of Ca^2+^ transients have strong frequency-dependent effects on DA release.^35, 47^ We found no difference in EC_50_ or Hill slope after 5-pulse 100hz stimulations (**Fig 4D**) (EC_50_: *Tor1a^+/+^*= 3.05 ± 2.45 mM; *Tor1a^+/ΔE^* = 1.49 ± 0.43 mM; Student’s *t*-test, p = 0.54, n=6; Hill coefficient: *Tor1a^+/ +^* = 1.16 ± 0.37; *Tor1a^+/ΔE^* = 1.94 ± 0.81; Student’s *t*-test, p = 0.40 n=6).

### Vesicle utilization in *Tor1a^+/ΔE^* mice

Snapin, which interacts with SNARE proteins, has been implicated as an interacting partner with torsinA.^48, 49^ Abnormalities in docking proteins can disrupt the vesicle refilling rate (i.e., the restoration of the readily releasable pool of vesicles after a release event) suggesting that the reduction in DA release in *Tor1a^+/ΔE^* knockin mice may result from sub-optimal vesicle utilization.^50^ To test this hypothesis, we conducted a rundown experiment by applying electrical stimulation at varying inter-pulse intervals (IPI) (5 min, 3 min, 1 min, 10 sec, 0.3 sec IPI) and examined the rate of decline in peak DA release. While long inter-stimulus intervals produced only moderate reductions in DA release (**Fig 5A-C**), DA release rapidly decreased at shorter inter-stimulus intervals of 10 s (**5D**) and 0.3 s intervals (**5E**; two-way ANOVA main effect of pulse number, F_3,30_ = 131.1, *p* < 0.0001). There was no difference between *Tor1a^+/+^* and *Tor1a^+/ΔE^* mice (two-way ANOVA main effect of genotype, F_1,10_ = 0.0017, p = 0.97) at the 0.3s interval. To obtain the rate of run-down, we plotted the DA released at the second pulse normalized to the first pulse at all stimulation intervals (**5F**). There was no difference in the rate of rundown between *Tor1a^+/+^* and *Tor1a^+/ΔE^* mice (two-way ANOVA genotype x IPI interaction effect, F_4,40_ = 0.766, p = 0.55).

**Fig 5:**
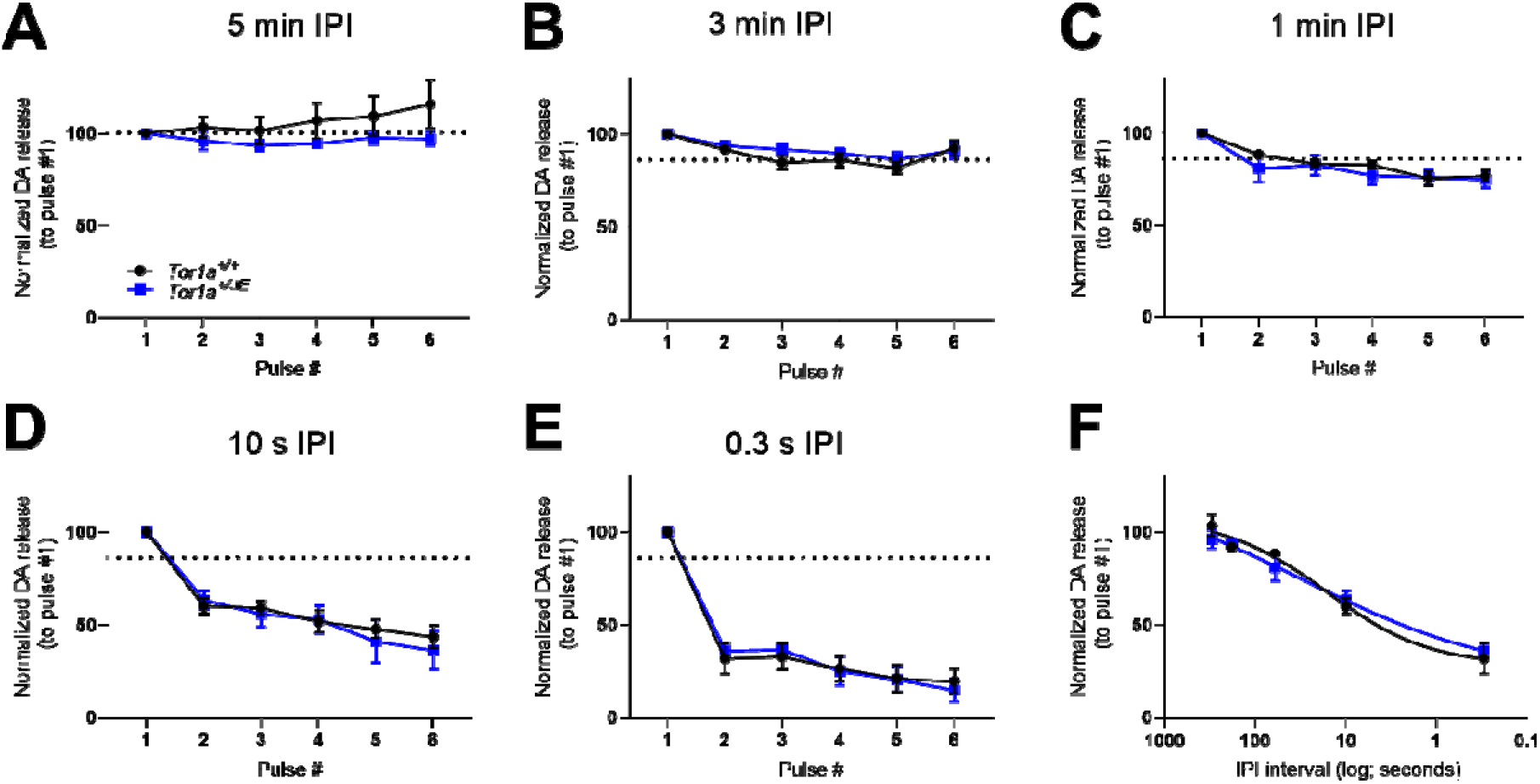
Rundown of evoked DA release after repeated simulation. DA release was recorded after sequential electrical stimulations with varying inter-pulse intervals (IPI) of 5 min (**A**), 3 min (**B**), 1 min (**C**), 10 sec (**D**), and 0.3 sec (**E**). (**F**) DA release at the second pulse normalized to the first pulse for all inter-pulse intervals (IPI) tested. There was no significant effect of genotype observed at any IPI (*n* = 6). Values represent mean ± SEM.

### D2 DA autoreceptor function in *Tor1a^+/ΔE^* mice

D2 DA autoreceptors located on the terminals of nigrostriatal neurons negatively regulate extracellular DA concentrations by mediating release.^51, 52^ A significant reduction in striatal D2 DA receptor availability is observed in human *DYT1-TOR1A* carriers but it is not known whether these defects are associated with autoreceptors, postsynaptic receptors or both.^53, 54^ To determine if abnormal D2 DA autoreceptor function mediates the deficit in DA release observed in *Tor1a^+/ΔE^* mice, we assessed the effect of the D2 DA receptor agonist quinpirole on DA release in *Tor1a^+/+^* and *Tor1a^+/ΔE^* mice (**Fig 6A**). Quinpirole dose-dependently decreased DA release in both genotypes, and there was no significant difference in IC_50_ between *Tor1a^+/+^* and *Tor1a^+/ΔE^* mice (IC_50_: *Tor1a^+/+^* = 87.59 ± 22.01 μM; *Tor1a^+/ΔE^* = 102.5 ± 39.97 μM; Student’s *t*-test, p= 0.43 n=5).

**Fig 6:**
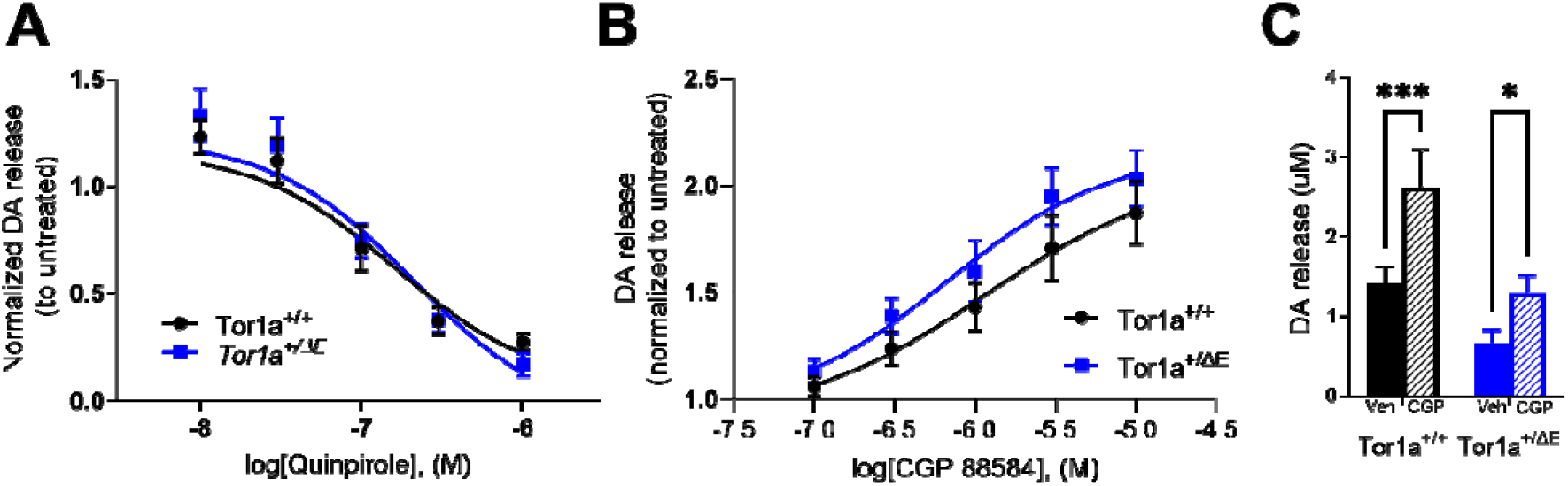
Sensitivity of *Tor1a^+/ΔE^* mice to D2 receptor activation or GABA_B_ receptor blockade. Control and *Tor1a^+/ΔE^* mice have similar sensitivity to the D2 DA receptor agonist quinpirole. There was no significant difference in IC_50_ between control and *Tor1a^+/ΔE^* mice (*n* = 5). (**B**) GABA_B_ receptor function in *Tor1a^+/ΔE^* mice. *Tor1a^+/+^* and *Tor1a^+/ΔE^* mice have simi**lar** sensitivity to the GABA_B_ receptor antagonist CGP 88584. There was no significant difference **in** EC_50_ between *Tor1a^+/+^* and *Tor1a^+/ΔE^* mice (*n* = 5). (**C**) Maximal effect of CGP 88584 (10 μM) on DA release. CGP 88584 significantly enhanced DA release in both genotypes, but there was no significant difference between the genotypes. Values represent mean ± SEM.

### GABA_B_ receptor function in *Tor1a^+/ΔE^* mice

Activation of GABA_B_ receptors on DA terminals reduces DA release in slice.^55^ Further, a recent study demonstrated that enhanced GABA tone in slice can mediate a ~50% reduction in DA release similar to that observed in *Tor1a^+/ΔE^* knockin mice.^56^ To determine if increased GABA_B_ receptor activation and/or increased GABA tone mediates the reduction in DA release in *Tor1a^+/ΔE^* knockin mice, we performed a dose response experiment with the selective GABA_B_ antagonist CGP 88584. CGP 88584 dose-dependently enhanced DA release in both *Tor1a^+/+^* and *Tor1a^+/ΔE^* mice (**Fig 6B**). However, there was no significant difference in either EC_50_ (EC_50_: *Tor1a^+/+^* = 1.16 ± 0.78 μM; *Tor1a^+/ΔE^* = 0.66 ± 0.39 μM; Student’s *t*-test, p = 0.58 n=5) or maximum effect between the genotypes at the highest drug concertation (10 μM CGP 88584) (main effect of treatment, two-way repeated measures ANOVA F_1,8_ = 36.24, p = 0.0003, genotype x treatment interaction effect, F_1,8_ = 3.85, p = 0.085) (**Fig 6C**).

### TorsinA localization in DA in neurons in *Tor1a^+/ΔE^* mice

The precise cellular localization of torsinA in neurons is controversial, with some studies demonstrating localization of torsinA at nerve terminals but others finding torsinA absent from nerve terminals.^57–59^ To begin to address if torsinA is localized to DA axons we performed immunohistochemistry experiments to determine torsinA localization in midbrain DA neurons and DA fibers coursing through the medial forebrain bundle. TorsinA immunofluorescence was observed in TH-positive soma but was not detected in the nearby TH-positive processes in both *Tor1a^+/+^* and *Tor1a^+/ΔE^* mice (**Fig 7A & B**). Although immunofluorescence for torsinA was observed in cell bodies adjacent to the medial forebrain bundle, it was not in TH-positive fibers within the medial forebrain bundle of either *Tor1a^+/+^* or *Tor1a^+/ΔE^* mice (**Fig 7C & D**).

**Fig 7.**
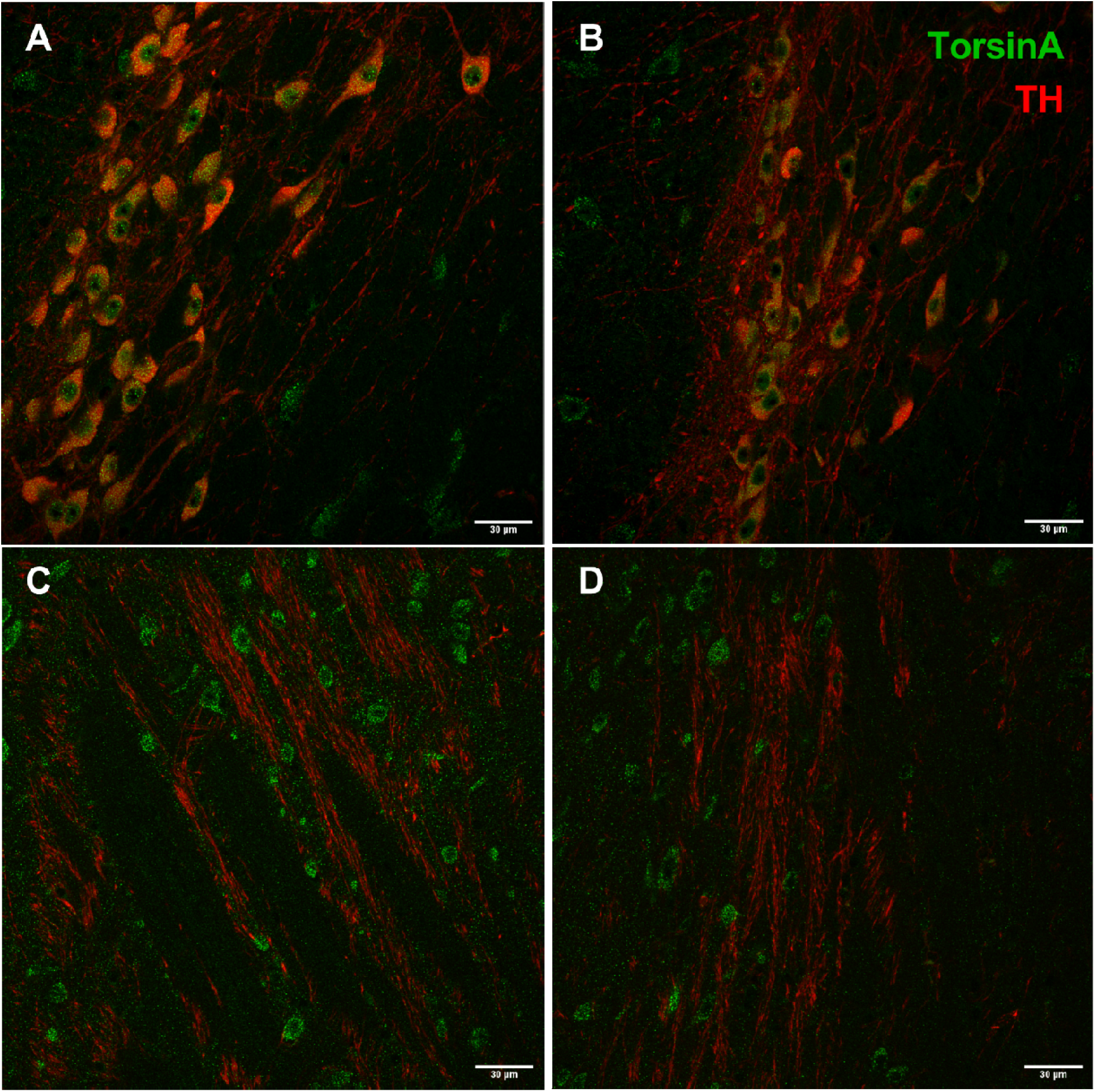
TorsinA and TH co-localization in DA neurons. Photomicrographs of SNc and medial forebrain bundle stained for TorsinA and TH in *Tor1a^+/+^* and *Tor1a^+/ΔE^* mice. TorsinA is dense**ly** localized to DA neuron cell bodies but absent from axons in both *Tor1a^+/+^* (**A**) and *Tor1a^+/ΔE^* mice (**B**). TorsinA is not localized to TH-positive fibers in the medial forebrain bundle in eith**er** *Tor1a^+/+^* (**C**) or *Tor1a^+/ΔE^* mice (**D**).

## DISCUSSION

Here, we demonstrate that the deficit in striatal DA release induced by *Tor1a(ΔE)* is caused by cell-intrinsic effects of the mutation within DA neurons while selective expression of *Tor1a(ΔE)* in striatal ChIs plays little, if any, role. Our results are in general agreement with a previous study demonstrating that transgenic overexpression of human torsinA(ΔE) in midbrain DA neurons causes a defect in DA release.^30^ However, in this same carefully-controlled study, overexpression of normal human torsinA also reduced striatal DA content, suggesting that overexpression, rather than the mutation *per se*, may have contributed to the abnormal dopaminergic response as it is known that overexpression of torsinA can produce extraneous CNS abnormalities that are unrelated to the mutation.^60^ Here, we expressed the human disease-causing genotype (*Tor1a^+/ΔE^*) in midbrain DA neurons via the endogenous gene, thus avoiding the confound of transgenic overexpression, to unambiguously pinpoint the site of dysfunction as DA neurons using both *in vitro* and *in vivo* approaches.

Although striatal cholinergic and dopaminergic signaling are inextricably linked, we found that selective expression of the disease-causing genotype (*Tor1a^+/ΔE^*) in cholinergic neurons did not influence DA release in the striatum. Our results are consistent with a previous study that found that selective knockout of *Tor1a* (*Tor1a^-/-^*) from cholinergic neurons does not affect the psychostimulant-induced increase in extracellular striatal DA.^61^ Taken together these findings are surprising because while cholinergic tone appears to be abnormal in *Tor1a^+/ΔE^* mice, it has little effect on DA neurotransmission.^11, 29^ It is possible that either the alteration in ACh tone is not due to direct action of *Tor1a(ΔE)* in ChIs and/or that abnormal cholinergic neurotransmission caused by the mutation is not sufficient to appreciably impact DA release.

Previous studies have demonstrated abnormal function of D2 DA receptors on ChIs of mice carrying the *Tor1a(ΔE)* mutation. Normally D2 DA receptor stimulation results in a decrease in firing rate of ChIs, but in mouse models of *DYT1-TOR1A* D2 DA receptor activation causes an increase in ChI firing rate.^10, 38, 62^ Current evidence suggests that this change in D2 DA receptor valence is due to hypercholinergia.^11^ However, it remains unclear if hypercholinergia itself is due to direct effects of *Tor1a(ΔE)* on ChIs or if it is an indirect effect of reduced DA release and this question will need to be addressed in future studies.

While our results demonstrate that intrinsic effects of *Tor1a(ΔE)* on DA neurons cause the deficit in DA release, the mechanism underlying this defect is unclear. Striatal tissue DA levels and the levels of DA metabolites in *DAT-cre; Tor1a^+/swap^* were normal which is in agreement with previous data in *Tor1a^+/ΔE^* knockin mice.^45^ Because our *ex vivo* brain slice preparation lacks DA neuron cell bodies, the mechanism leading to reduced DA release cannot be explained by firing rates. Although the localization of torsinA within neurons is debated, ^57–59^ our results suggest that torsinA is predominantly localized to DA neuron cell bodies as torsinA was not detected in TH-positive neuronal processes, consistent with the localization of torsinA and its activators LAP1 and LULL1 to the nuclear membrane and endoplasmic reticulum.^63, 64^ While we have not conclusively ruled out the presence of torsinA in axons, it is reasonable to postulate that the *Tor1a(ΔE)*-induced reduction in DA release does not result from direct effects of the mutant protein at the presynaptic terminal *per se* considering that, with the exception of the robust reduction in the amount of DA released, DA terminal function was largely intact.

Our data does offer clues to potential mechanisms that mediate the *Tor1a(ΔE)*-induced reduction in striatal DA release. *Tor1a(ΔE)* may disrupt DA release indirectly by disrupting basic cellular machinery in DA cell bodies. *Tor1a* has been linked with a variety of different cellular process, including: membrane protein trafficking, secretory protein processing, endoplasmic reticulum stress response, lipid metabolism, nuclear membrane function and maintenance, and many others.^64–73^ The strong link between *Tor1a* and endoplasmic reticulum function suggests the hypothesis that mutant *Tor1a(ΔE)* disrupts posttranslational processing of synaptic proteins or disrupts the lipid constitution of vesicles and/or cell membrane to reduce the probability of release. Alternatively, *Tor1a(ΔE)* may disrupt the normal development of DA neurons. For example, *Tor1a(ΔE)* may result in a decrease in the number of DA release sites in the striatum. While the cellular processes that regulate the number of DA release sites are poorly understood, it is possible that *Tor1a(ΔE)* reduces the number of striatal DA release sites by modulating glia-derived neurotrophic factor GDNF, which is known to enhance striatal DA release sites.^74^

Taken together, our data demonstrate that the reduction in striatal DA release observed in mouse models of *DYT1-TOR1A* dystonia is due to intrinsic effects of *Tor1a(ΔE)* on midbrain DA neurons and not ChIs. While the precise mechanism linking *Tor1a(ΔE)* to reduced DA release remains unknown, these results point to DA neurons as a cellular target for new therapeutics.

## ACKNOWLEDGMENTS

This research project was supported in part by the Emory University Integrated Cellular Imaging Core and Emory HPLC Bioanalytical Core.

## Abbreviations

ACh: acetylcholine
aCSF: artificial cerebrospinal fluid
BSA: bovine serum albumin
ChAT: choline acetyltransferase
ChI: cholinergic interneuron
DA: dopamine
DEPC: diethyl pyrocarboante
FSCV: fast scan cyclic voltammetry
HPLC: high performance liquid chromatography
IPI: inter-pulse interval
LCM: laser capture microdissection
PBS: phosphate-buffered saline
SNc: substantia nigra pars compacta
TBS: tris-buffered saline
TH: tyrosine hydroxylase
VTA: ventral tegmental area

